# Capillary connections between sensory circumventricular organs and adjacent parenchyma enable local volume transmission

**DOI:** 10.1101/2024.07.30.605849

**Authors:** Yifan Yao, Yannan Chen, Raju Tomer, Rae Silver

## Abstract

Among contributors to diffusible signaling are portal systems which join two capillary beds through connecting veins (Dorland 2020). Portal systems allow diffusible signals to be transported in high concentrations directly from one capillary bed to the other without dilution in the systemic circulation. Two portal systems have been identified in the brain. The first was discovered almost a century ago and connects the median eminence to the anterior pituitary gland (Popa & Fielding 1930). The second was discovered a few years ago, and links the suprachiasmatic nucleus to the organum vasculosum of the lamina terminalis, a sensory circumventricular organ (CVO) (Yao et al. 2021). Sensory CVOs bear neuronal receptors for sensing signals in the fluid milieu (McKinley et al. 2003). They line the surface of brain ventricles and bear fenestrated capillaries, thereby lacking blood brain barriers. It is not known whether the other sensory CVOs, namely the subfornical organ (SFO), and area postrema (AP) form portal neurovascular connections with nearby parenchymal tissue. This has been difficult to establish as the structures lie at the midline and protrude into the ventricular space. To preserve the integrity of the vasculature of CVOs and their adjacent neuropil, we combined iDISCO clearing and light-sheet microscopy to acquire volumetric images of blood vessels. The results indicate that there is a portal pathway linking the capillary vessels of the SFO and the posterior septal nuclei, namely the septofimbrial nucleus and the triangular nucleus of the septum. Unlike the latter arrangement, the AP and the nucleus of the solitary tract share their capillary beds. Taken together, the results reveal that all three sensory circumventricular organs bear specialized capillary connections to adjacent neuropil, providing a direct route for diffusible signals to travel from their source to their targets.

## 1 INTRODUCTION

With the increased understanding of volume transmission (VT) mechanisms^1^, the discovery of the glymphatic system^2^, and the distinct role of chemical secretions in modulating neural wiring networks^3^, it is clear that fluid systems of the brain, including the cerebrospinal fluid and perivascular spaces are key players in central nervous system function and dysfunction^4,5^. Among the important elements are circumventricular organs (CVOs), a group of structures lining the surface of the ventricles. They have large perivascular spaces, are in contact with cerebrospinal fluid (CSF), are highly vascularized, and most feature fenestrated capillaries, as reviewed in^6^. These properties have led to the concept of CVOs as “windows of the brains”^7^. CVOs are categorized into two groups – secretory and sensory^6^. The sensory CVOs include the organum vasculosum of the lamina terminalis (OVLT), subfornical organ (SFO), and area postrema (AP). They contain neurons expressing a wide variety of receptors to blood-borne signals, CSF contents, or neural inputs to relay sensed changes in fluid milieu to other regions of the brain, further modulating behavior and physiology^8,9^.

It is now known that there is a venous portal pathway connecting the capillary beds of the suprachiasmatic nucleus (SCN) to those of the OVLT^10,11^. However, whether each of the other sensory CVOs connects to the parenchyma, to which they are attached via a portal pathway, remains unclear. In the literature on the vascular anatomy of sensory CVOs, there are suggestions of joined capillary beds between the other sensory CVO’s and their respective adjacent parenchyma. Thus, Akert in 1969^12^ commented on densely packed SFO capillary loops, and wrote, “It seems that the next step would consist in the intravital study of blood flow with special attention to the question of a portal circulation in the direction of the certain hypothalamic and septal areas.” Roth and Yamamoto in 1968^13^ also reported “two distinctively different capillary beds, one in the AP and the other in the medulla, are seen to be joined by short interconnecting vessels”. The anatomical relationship between the SFO, AP and adjacent parenchyma has been described in several species, including mouse (reviewed in ^14-16^). The SFO is located along the anterior-dorsal wall of the third ventricle (3V), where it lies at the dorsal extremity of the lamina terminalis. It is apposed to the ventral hippocampal commissure (VHC), is separated from the septal area by the VHC and extends into the lumen of the 3V where it lies adjacent to the choroid plexus (CP) near the ceiling of the 3V^14^. The AP resides on the floor of the fourth ventricle (4V) and is attached to the nucleus of the solitary tract (NTS) at its ventral surface.

Portal systems in the brain are key to integrating brain and bodily functions. They are vascular arrangement where two capillary beds are linked by connecting veins^17^. Portal pathways enable blood-borne substances to be transported in high concentrations from the capillary bed of one region to the capillary bed of a local target site without dilution in the systemic blood supply^17^. The pituitary portal system was first identified almost 100 years ago^18,19^. Here, portal veins connect the capillary beds of the median eminence (ME) to those of the anterior pituitary gland. Hypothalamic-releasing hormones from the ME travel in the portal veins lying in the pituitary stalk to reach the capillary bed of the anterior pituitary gland. In the anterior pituitary gland, hypothalamic-releasing hormones stimulate the production of the pituitary trophic hormones, including the adrenocorticotropic hormone, thyroid-stimulating hormone, luteinizing hormone and follicle-stimulating hormone^20^. For example, the portal pathway enables a small population of about 600 GnRH neurons are sufficient to support reproductive responses^21^. In this way, the pituitary portal system, via the pituitary gland, serves as a relay, amplifying signals from a small population of hypothalamic neurons to impact body-wide functions necessary for survival, including stress responses, reproduction, and metabolism^22-25^. The SCN, like the pituitary gland, has body-wide impact^26,27^. This small nucleus serves as a daily clock which sets the phase of rhythms in “clock cells” throughout the body^28-30^. The SCN has both neural and vascular-portal connections to the OVLT^10,31^ and the OVLT in turn, has extensive neural and vascular connections to the rest of the brain^6,32,33^. The foregoing evidence highlights the possibility that sensory CVOs are positioned to amplify neurosecretory signals from small populations of hypothalamic neurons.

Though the vasculature of SFO and AP have been described previously, capillary blood vessels and connections to adjacent parenchyma and choroid plexus (ChP) have not been examined. Our goal was to examine the capillary vasculature organization of the SFO and AP in relation to the parenchymal tissue and the ChP to which they are attached using a preparation that enables studying intact blood vessels (BVs) and does not require tissue. We scanned iDISCO-cleared whole mouse brain vasculature using light sheet microscopy and immunochemistry for collagen to label the entire vasculature. We then conducted brain registration aligned to the Allen Mouse Brain Atlas (ABA) and computer-aided BV tracing to localize the capillary connections between SFO and AP and their adjacent brain regions, respectively. To test the findings more directly, we next examined the capillary organization of the regions that had been identified in the registered results in a separate series, using brain region-specific markers to definitively delineate SFO, AP, their adjacent parenchyma and their capillary networks.

## 2 METHODS

### 2.1 Terminology

We adopted the nomenclature of ABA^34^ (Allen Institute for Brain Science, 2004) or Paxinos and Franklin’s the Mouse Brain in Stereotaxic Coordinates^35^ in describing the brain regions studied herein.

### 2.2 Animals and housing

Male C3H mice (n=4) and C57BL/6J mice (n=4) aged 8–10 weeks were purchased from the Jackson Laboratory (Sacramento, CA). C3H mice were used for whole brain imaging and vasculature registration while C57BL/6J were used for brain slices for labeling specific nuclei. C3H mice produce melatonin in the pineal gland^36^, while the more commonly used C57BL/6J animals do not. The mice were housed in the lab for at least two weeks after arrival prior to being used in these studies. All animals were provided with ad libitum access to food and water, and maintained in a 12:12-h light:dark (LD) cycle (lights-on at 7:00 AM, where zeitgeber time (ZT) 0 refers to the time of lights on). All experiments were carried out in accordance with the guidelines of Columbia University’s Institutional Animal Care and Use Committee protocol number AC-AABH1603.

### 2.3 Perfusion

At ZT6 animals were deeply anesthetized (11%ketamine+2%xylazine, 10ml/kg body weight i.p.) and perfused intracardially with 50 ml 0.9% saline followed by 100 ml 4% paraformaldehyde (PFA) in 0.1 M phosphate buffer (PB), pH 7.3. Brains were post-fixed at 4 °C overnight. The next day, brains for iDISCO were washed with PBS and stored in PBS with 0.02% sodium azide at 4°C until future use.

### 2.4 Brain clearing and immunostaining for whole brain and brain slices

The experimental pipeline is shown in Figure 1. For both whole brains of C3H mouse and brain slices of C57BL/6J mouse, the iDISCO protocol was used (Renier et al., 2016) with the following modifications. For whole brain analysis, primary antibody goat anti-type IV Collagen-UNLB (1:50, SouthernBiotech, Birmingham, AL) was used to label the entire vasculature and secondary antibody donkey anti-goat Cy5 (1:200, Jackson ImmunoResearch) was used to identify collagen. Brains were incubated with the primary antibody for two weeks followed by secondary antibody for one week.

**Figure 1:**
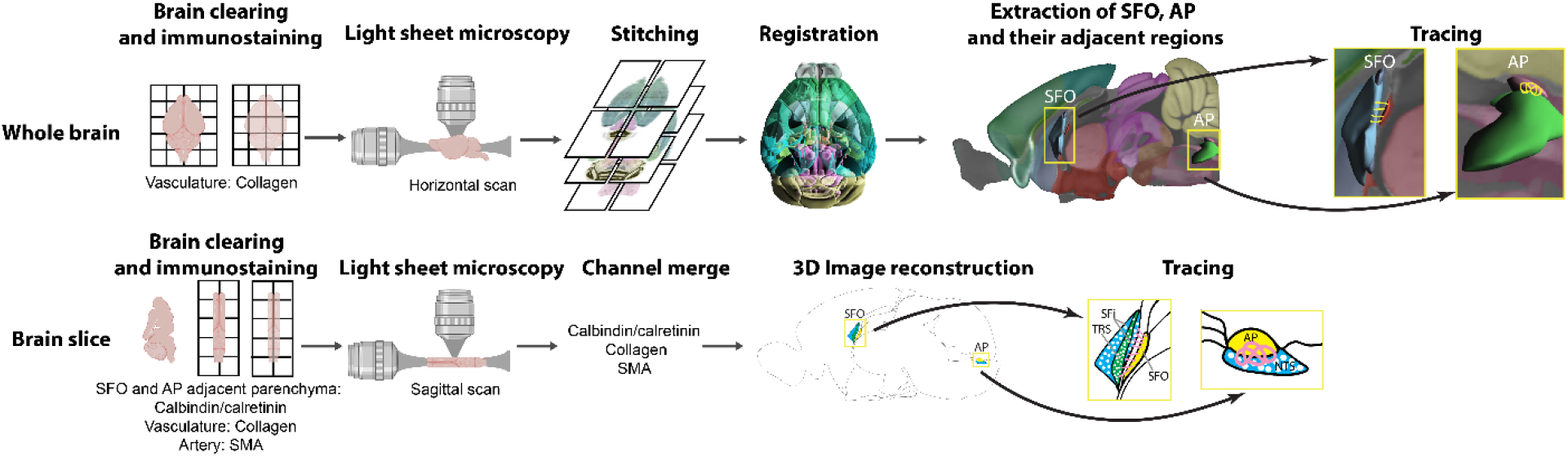
Experimental pipeline. Top row, procedure for whole brain clearing, imaging and vascular connection analysis. Bottom row, procedure for brain slice (4mm) clearing, imaging and vascular connection analysis. Abbreviations: AP=area postrema; NTS=nucleus of solitary tract; SFi=septofimbrial nucleus; SFO=subfornical organ; SMA=smooth muscle actin; TRS=triangular nucleus of septum.

For the brain slices, 2mm of tissue on each side of the midline in the sagittal plane were kept with a steel brain matrix to preserve midline CVOs and their nearby regions. The primary antibody solution contains mouse anti-smooth muscle actin (SMA) antibody (1:67, Dako, M0851, Santa Clara, CA) for labeling arteries, goat anti-collagen (as above) for labeling the entire vasculature, and a mixed solution of rabbit anti-calretinin (1:500, Chemicon, AB149, Temecula, CA) and rabbit anti-calbindin (1:5000, Swant, CB38a, Burgdorf, Switzerland) to identify regions adjacent to the SFO and AP^37,38^. The secondary antibody donkey anti-rabbit Cy2 (1:200, Jackson ImmunoResearch) was used to detect calretinin and calbindin. Donkey anti-mouse Cy3 (1:200, Jackson ImmunoResearch) was used to detect SMA. Donkey anti-goat Cy5 (1:200, Jackson ImmunoResearch) was for detecting collagen. Tissues were incubated with primary antibodies for one week and then with secondary antibodies for four days.

### 2.5 Light-sheet microscopy

Whole brain samples were imaged using CLARITY-Optimized Light Sheet Microscopy (COLM) as previously described^39^. An Olympus 10x objective was used and achieved a pixel size of 0.585µm, with further downsampling applied for efficient processing. The image tiles were stitched using BigStitcher^40^ with careful manual inspection to ensure vasculature connectivity.

For brain slices, images were acquired with Ultramicroscope II (LaVision BioTec, Bielefeld, Germany) equipped with an Olympus MVX10 zoom body (Olympus, Barlett, TN), a LaVision BioTec Laser Module (LaVision BioTec), and an Andor Neo sCMOS Camera (Andor Technology, Concord, MA) with a pixel size of 6.5 µm. The following lasers and filters were used: Cy2-calretinin/calbindin was excited at 488nm and emission was acquired at 525±50nm; Cy3-SMA was excited at 561nm and acquired by 605±50nm; Cy5-collagen was excited at 639nm and emission was collected at 705±72nm. The vasculature of the SFO and the AP regions were scanned horizontally with a voxel size 0.94 × 0.94 × 1µm (left-right x rostral-caudal x dorsal-ventral).

### 2.6 Whole brain registration

The whole brain samples were aligned using suiteWB pipeline^41^ as previously described. Specifically, suiteWB employs a multi-step multi-resolution 3D registration approach using Mutual Information as the similarity metric. The registration process initiates with a rigid transformation, incorporating six degrees of freedom for rotation and translation, aligning the moving image (i.e., the image being aligned) roughly with the reference image. Subsequently, the moving image undergoes an affine transformation involving twelve degrees of freedom to address shearing and shrinking artifacts introduced during labeling, tissue clearing, and imaging. Finally, a uniform grid of control points and third-order B-splines are used to account for non-rigid local transformations. Affine and nonrigid steps were executed at three different resolutions. The algorithms were implemented with ITKv4^42^. All samples underwent registration using 488nm channel onto our local average reference brain at 10µm resolution with corresponding ABA brain region annotations, and the resultant spatial transformation parameters were applied to high-resolution datasets with lateral resolution 2.5µm and axial resolution 5µm.

### 2.7 Extraction of CVOs and neighboring regions

CVOs were initially identified using the ABA brain region annotations, and neighboring regions were chosen based on the 8th level of the ABA hierarchy tree, encompassing areas within a 25µm distance of the CVO bounding box. Subsequently, a cubic Region of Interest (ROI) with edges measuring 1250µm was cropped, centered on the CVO, to facilitate detailed manual inspection and tracing. The alignment between the ABA annotation mask and the brain within the ROI was further refined manually through translation and rotation adjustments.

### 2.8 Tracing

Acquired samples were imported into Imaris (Bitplane AG, Zurich, Switzerland) for tracing. Tracing was done with the “filament” model in the collagen-labelled channel. The autodepth mode was used with auto-center and auto-diameter correction.

### 2.9 BV diameter measurement and statistical analysis

The flowing types of BV segments were measured: SFO capillary, SFO portal vessels, septofimbrial nucleus and triangular nucleus of septum capillaries, AP capillaries and nucleus of the solitary tract capillaries. 20 of each type of BV segments were randomly chosen from light sheet images of brain slices of C57BL/6J mice and measured with the “Straight” line tool in Fiji ImageJ^43^. The capillaries are identified as BV segments that have diameters below 10µm and have no SMA staining. SFO portal vessels are identified by its location as the intervening vessel between the two capillary beds of SFO and the adjacent posterior septal nuclei. Capillaries of AP and its neighboring NTS are differentiated by distinct collagen staining intensity in either region, where AP has stronger staining than NTS (original measurements are documented in Supplementary table 1). All groups of diameter measurements are normally distributed and establish sphericity (Supplementary table 2). All statistical analyses were done with Python 3.0 and its packages NumPy^44^, Pandas^45^ and Pingouin^46^. Graphs were created with Python packages Matplotlib^47^ and Seaborn^48^.

## 3 RESULTS

### 3.1 Vascular connections between SFO and adjacent parenchyma in iDISCO cleared, registered brain

The rostral SFO lies near the triangular nucleus of the septum (TRS) and septofimbrial nucleus, but is separated from these nuclei by the ventromedial hippocampal commissure (VHC, Figure 2A-D). The TRS and SFi together, have been described as the posterior septal nuclei^49,50^. As seen in the collagen labelled vasculature registered to the ABA, the dorsal, caudal, and ventral aspects of the SFO protrude into the 3V (Figure 2E, sagittal view). Our whole brain registration material suggests that the rostral SFO is attached to the VHC whose upper 4/5^th^ abuts the SFI and its lower 1/5^th^ abuts the TRS.

**Figure 2:**
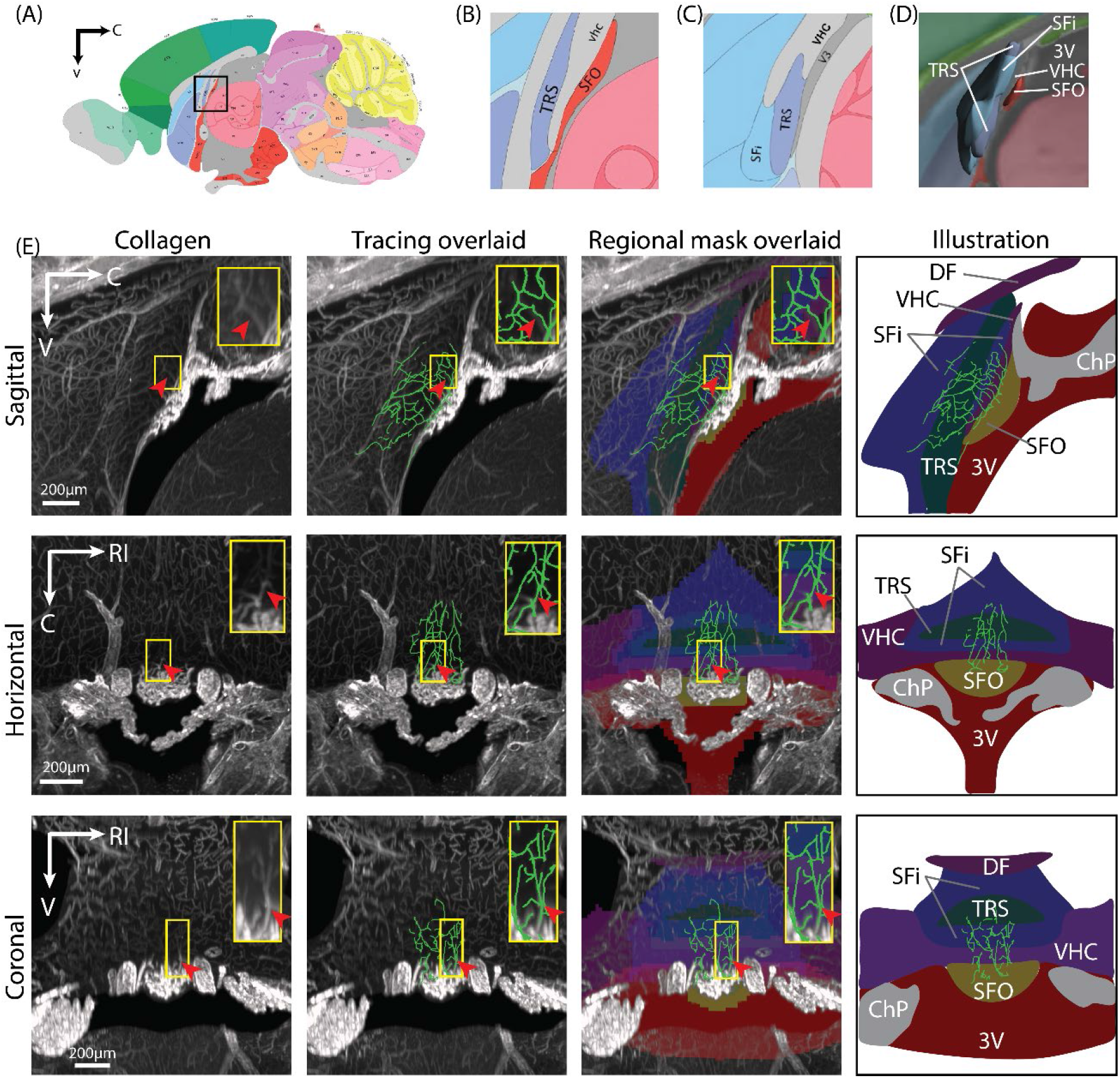
Vasculature of SFO and Adjacent Parenchymal Region using the ABA for Registration. (A) Schematic of the brain (P56 sagittal view #20, <u>Allen</u> Institute for Brain Science, 2004) in sagittal view; the boxed area demarcates the region containing the SFO. (B) Magnified view of the boxed area of (A). (C) Magnified view of region lateral to b showing the spatial relationship between TRS and SFi (P56 sagittal view #17 in ABA). SFO is not present at locus. (D) 3D atlas view (ABA) enabling a view of SFi, TRS, VHC and 3V, all regions adjacent to the SFO. (E) Vascular connections between SFO and the SFi and TRS regions in sagittal (upper row), horizontal (middle row) and coronal (lower row) orientations. Depicted from left to right: collagen labeling of the vasculature of the SFO and its nearby regions, tracing overlaid on vasculature staining, registered regional masks merged with traces and vasculature, and a summary illustration. Insets highlight vascular connections (arrowheads) between the SFO and the posterior septal nuclei (which encompasses both SFi and TRS). z = 100μm. Color code: ChP=gray; DF and VHC=dark purple; SFi=dark blue; SFO=dark yellow; TRS=dark green; 3V=dark red. Abbreviations: ABA=Allen mouse brain atlas; C=caudal; ChP=choroid plexus; DF=dorsal fornix; LE=left; RI=right; V=ventral; VHC=ventral hippocampal commissure; 3V=third ventricle. Remaining abbreviations as in Figure 1.

In the SFO, shown in sagittal, horizontal and coronal orientations, BVs identified by collagen labeling form an intensely tangled web that protrudes into the 3V (Figure 2E, left panel). In contrast, the BVs lying rostral to the SFO, encompassing the posterior septal nuclei, are sparser. In the sagittal view of SFO, the collagen-labelled and the traced BVs indicate that several vessel segments emerge from SFO capillaries along its entire dorsoventral axis (Figure 2E, top row). In the upper 4/5^th^ where the VHC lies between the SFO and SFi, the BV segments penetrate the VHC, and first arborize into capillaries in the SFi and then further arborize in the TRS. In the lower 1/5^th^ of the SFO, where the ventral VHC directly contacts the TRS, the BV segments emerging from the bottom of the SFO directly enter the TRS (Figure 2E, top row). In the horizontal and the coronal view (Figure 2E, middle & bottom row), the vascular connection between the SFO capillaries and those in the posterior septal nuclei can also be seen along the mediolateral axes of these nuclei. It remains to be determined whether these are connecting venules or direct capillary connections. It also remains to be proven that the registered material correctly identifies the brain regions. In order to further characterize the BV connections between the SFO and the posterior septal nuclei, we next identified the SFi and TRS using immunostaining.

### 3.2 Vascular connections between SFO and adjacent parenchyma in iDISCO cleared immunostained brain sections

The immunostaining results indicate that the size of calbindin/calretinin stained neurons in SFi are larger than those in TRS, consistent with previous reports that the SFi contains “septal giant cells” (15-21 um, scattered), whereas those in the TRS have a relatively smaller size (7-14 um, densely packed^49 51^. The localization of these nuclei confirms the results seen in brains registered to the ABA showing that VHC lies between the SFO and the posterior septal nuclei (Figure 3A). Importantly, several BV segments emerge from the SFO along its rostral surface, traverse the VHC (Figure 3B). At its upper 4/5^th^ of the VHC attached to SFO, the portal vessels branch to join the capillaries of SFi (Figure 3C). At the lower 1/5^th^, the portal vessels first travel ventrally, and then arborize and connect to the capillaries of the TRS. and connect to the capillaries of the SFi and TRS (Figure 3D). The trunks of these portal venules are BV segments that bear no branch points. As is typical of portal venules, the diameter of the portal venules penetrating the VHC are significantly larger than the capillaries in either the SFO or the posterior septal nuclei (Figure 3E).

**Figure 3:**
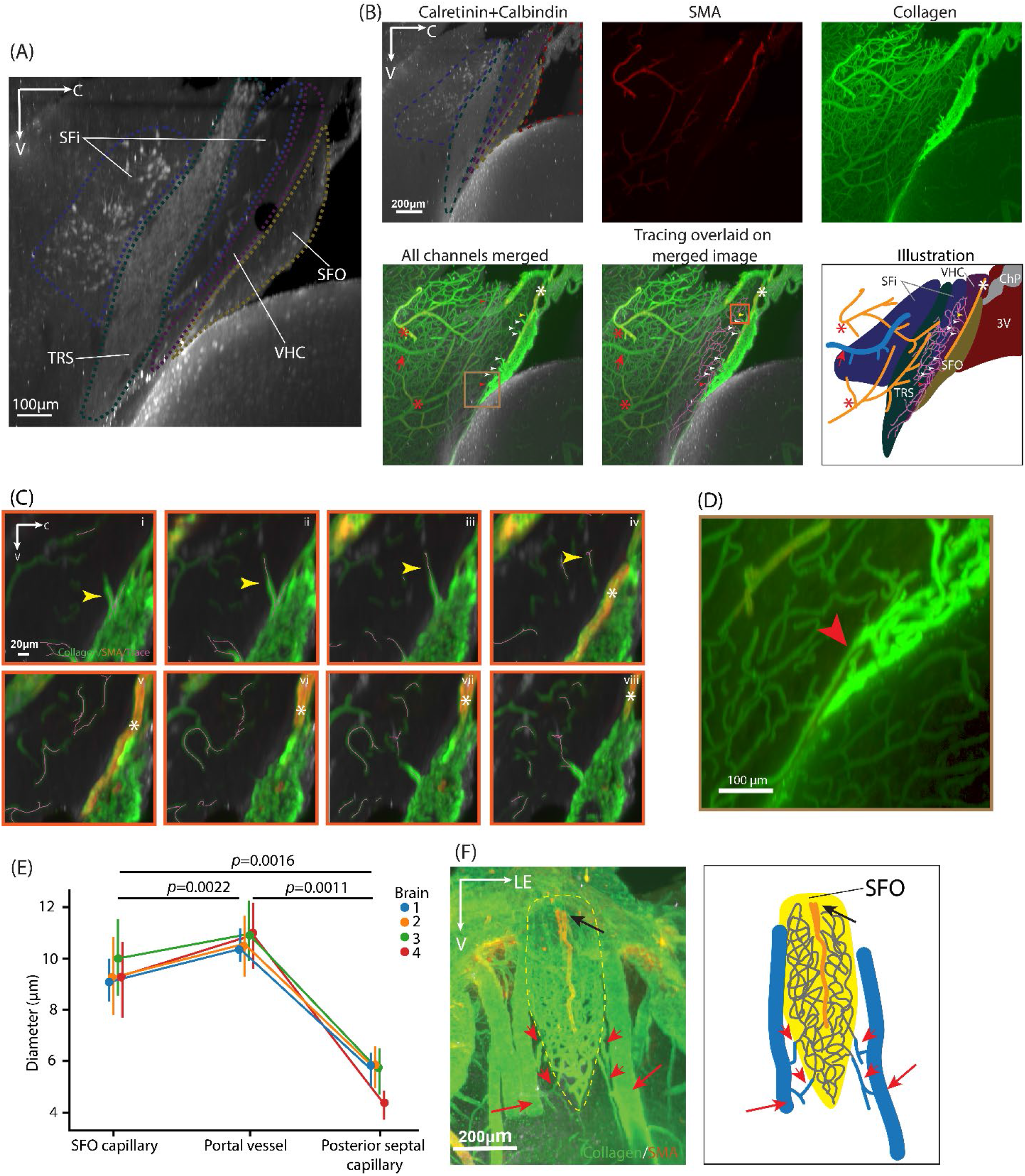
Vascular Connections between SFO to TRS and SFi in calbindin/ calretinin immunostained sections. (A) The full extent of the SFO abuts the VHC. At its upper 4/5th the VHC abuts the SFi while at its lower 1/5^th^ it abuts the TRS. The SFi wraps around TRS. (B) Vasculature of SFO, SFi and TRS in sagittal orientation. From left to right, the upper row shows immunochemical delineation of the SFi and TRS region, and the bottom row merges the immunostained channels followed by panels showing blood vessel tracing, and an illustration summarizing all these elements. Portal vessels (arrowheads) at the rostral aspect of the SFO penetrate the VHC. and join the capillary vessels in the SFi and in the TRS. The artery supplying the SFO (white asterisk) enters from its dorsal aspect. Arteries supplying the posterior septal nuclei enter at the rostrodorsal and rostroventral aspects (red asterisk). The draining vein of the posterior septal nuclei leaves from its rostrodorsal aspect (red arrow). (C) Magnified views of a portal vessel highlighted by the yellow arrowhead in the orange boxed area in the tracing overlaid on the merged image panel (C). (i-iv) A portal vessel (yellow arrowhead) penetrates the VHC and connects the rostral SFO capillaries to those of the SFi, shown in serial sections in the z axis. (iv-viii) An artery (asterisk) supplying SFO lies above the point of emergence of the portal vessel, shown in serial sections in the z axis. The artery lies ∼12µm above the portal vessel (plate iii vs iv). (D) Magnified the view of a portal vessel connects SFO and TRS highlighted by the red arrowhead in the brown boxed area in all channels merged panel in (B). A portal vessel (red arrowhead) first travels ventrally before branching and joining the capillaries in the TRS. (E) Diameters 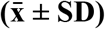 of SFO and posterior septal capillaries and connecting portal vessels (4 brains, 5 BV segments of each category of each brain) were as follows: SFO (9.42±1.61 µm), portal vessels (10.70±1.31 µm), and posterior septum (5.33±1.02µm). Statistical analysis was performed using repeated measurement ANOVA (F(2,6) = 140.2720, p=9.1808×10^−6^) followed by post-hoc multiple pairwise comparisons with Benjamini/Hochberg FDR correction: SFO capillary versus portal vessel (uncorrected p=0.002229, corrected p = 0.002229); SFO capillary versus posterior septal capillary (uncorrected p = 0.001598, corrected p = 0.002229); portal vessel versus posterior septal capillary (uncorrected p = 0.001081, corrected p = 0.002229). Detailed statistical results are in Supplementary Table 2. Data in the plot shown as 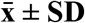. (F) Arterial supply and venous drainage of SFO in coronal orientation shows immunostaining of SFO vasculature (left) and corresponding illustration (right). The supplying artery travels along the dorsoventral axis (black arrow) of the SFO, extending 2/3 of its length. Small veins form near the lateral edges of both sides of the SFO (red arrowheads), and then drain to larger veins (red arrows). In (A, C) z = 150μm; in (B) z =12μm; in (E) z = 300μm. Color code: arteries=orange; capillaries=gray; yellow=SFO; veins=blue. Abbreviations as in Figure 1, 2.

### 3.3 Arterial supply and venous drainage of SFO and the posterior septal nuclei

The arteries and veins of the SFO and the posterior septal nuclei are clearly seen in the calbindin/calretinin labeled sample. In the sagittal view, the arterial supply of SFO enters its dorsal aspect and travels ventrally along the rostral surface of SFO, at the interface between SFO and the VHC (Figure 3B). In coronal view, the SFO artery travels along the midline, running dorsoventrally (Figure 3F), extending 2/3^rd^ of the length of the dorsoventral axis of SFO. At its bottom 1/3^rd^, the SFO artery arborizes into arterioles and capillaries. The arteries supplying the posterior septal nuclei have two sources, seen in sagittal view. One enters the posterior septal region from its rostroventral aspect. The branches of this artery mostly extend caudally and vascularize the part of posterior septal neuropils closely attached to the VHC. The second artery supplying the posterior septal nuclei enters this region from its rostral aspect. Some arteriole branches of this artery arborize and then join the capillaries in the caudal part of the posterior septal nuclei. As seen in the coronal view, the draining veins of the SFO are a series of small veins formed on the lateral edges along its dorsoventral axis. They run either laterally or ventrally and then merge into the larger vein next to the SFO (Figure 3F). In sagittal view, it can be seen that the draining vein of the posterior septal nuclei collects the capillaries from the rostral region, in close apposition to its supplying artery (Figure 3B).

### 3.4 Vascular connections between AP and adjacent parenchyma in iDISCO cleared, registered brain

We next examined AP and its neighboring regions in the registered sample (Figure 4A). As in prior work in several species^52^, we find that the AP is located at the caudal end of the 4V floor (Figure 4A,B). Its rostral aspect protrudes into the CSF. The ventral AP is in close contact with NTS, and the major volume of the AP is embedded in the latter. Of interest here is the capillary organization of the AP and NTS (Figure 4C). The AP and NTS capillaries can be distinguished by differences in the intensity of collagen staining, with much stronger collagen expression in AP BVs. In the sagittal view of the medial plane of the brain (Figure 4C, top row), the ventrocaudal part of the AP capillaries spread into the NTS. At its most rostral tip (arrow) capillaries of the AP are connected to the ChP in the 4V. In the horizontal and coronal view (Figure 4C, middle & bottom row), the capillaries of lateral AP merge into the surrounding NTS and the rostral AP is in contact with the wall of 4V. In the coronal view, capillaries in the ventrolateral aspect of AP form web-like connection with the adjacent NTS. Tracing of the AP capillaries shows that they extend into ventral, lateral, caudal directions and are continuous with the capillaries of the NTS (Figure 4C, regional mask overlaid on collagen labelled and traced vessels).

**Figure 4:**
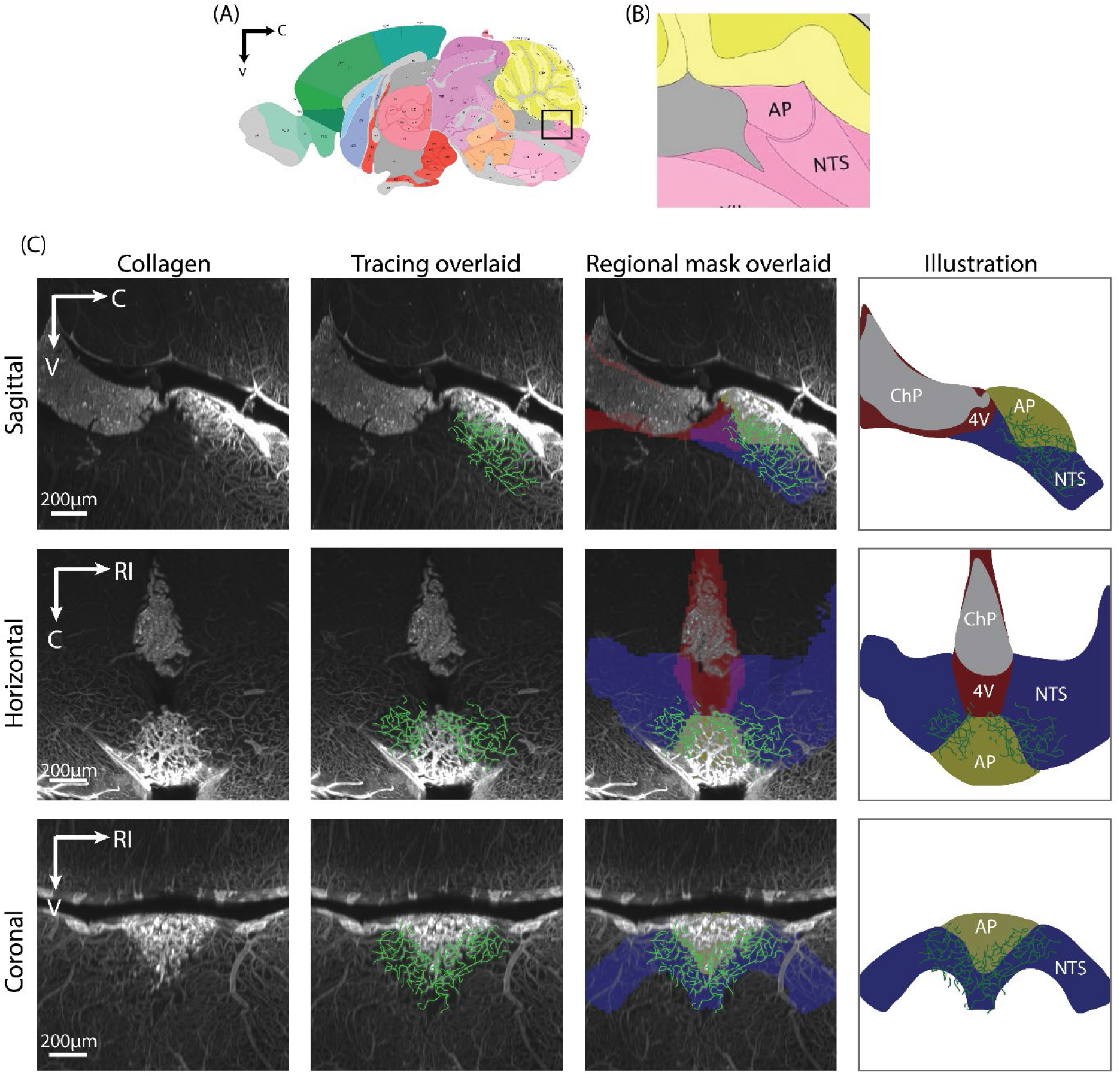
Identification of AP and Adjacent Parenchymal Vasculature using the ABA for Registration. (A) Schematic of the brain and the boxed area demarcating the region containing AP in the sagittal view. (Schematic from ABA). (B) Magnified view of the boxed region of (A). (C) Vascular connections between AP and NTS in sagittal (upper row), horizontal (middle row) and coronal (lower row) orientations. Depicted from left to right panels: collagen labeling of the AP and its nearby vasculature, tracing overlaid on blood vessel staining, registered regional masks overlaid with traces and blood vessels staining, and a summary illustration. The shared capillary network of AP and NTS can be seen in all three orientations. z=150μm. Color code: AP=yellow; ChP=gray; NTS=blue; 4V=red. Abbreviation: 4V=fourth ventricle. The remaining color codes and abbreviations as in Figures 1-3.

Next, calbindin and calretinin antibodies were used to definitively identify the NTS^37^ in order to evaluate the precise relationships of the capillary vessels of the AP and NTS in this preparation. Calbindin/calretinin positive cells are seen in the region adjacent to the AP in sagittal, horizontal and coronal orientations (Figure 5A). The results indicate that the capillary vessels from the ventral surface of the AP extend into the NTS and merge with the NTS capillaries, seen in a high power view at the interface of these regions (Figure 5B). The capillaries from both regions form a fully intertwined capillary network. The more intense collagen signal in the AP capillaries compared to those of the NTS is due to the different sizes of capillaries in these two regions, with larger capillaries in the AP than in the NTS (Figure 5C).

**Figure 5:**
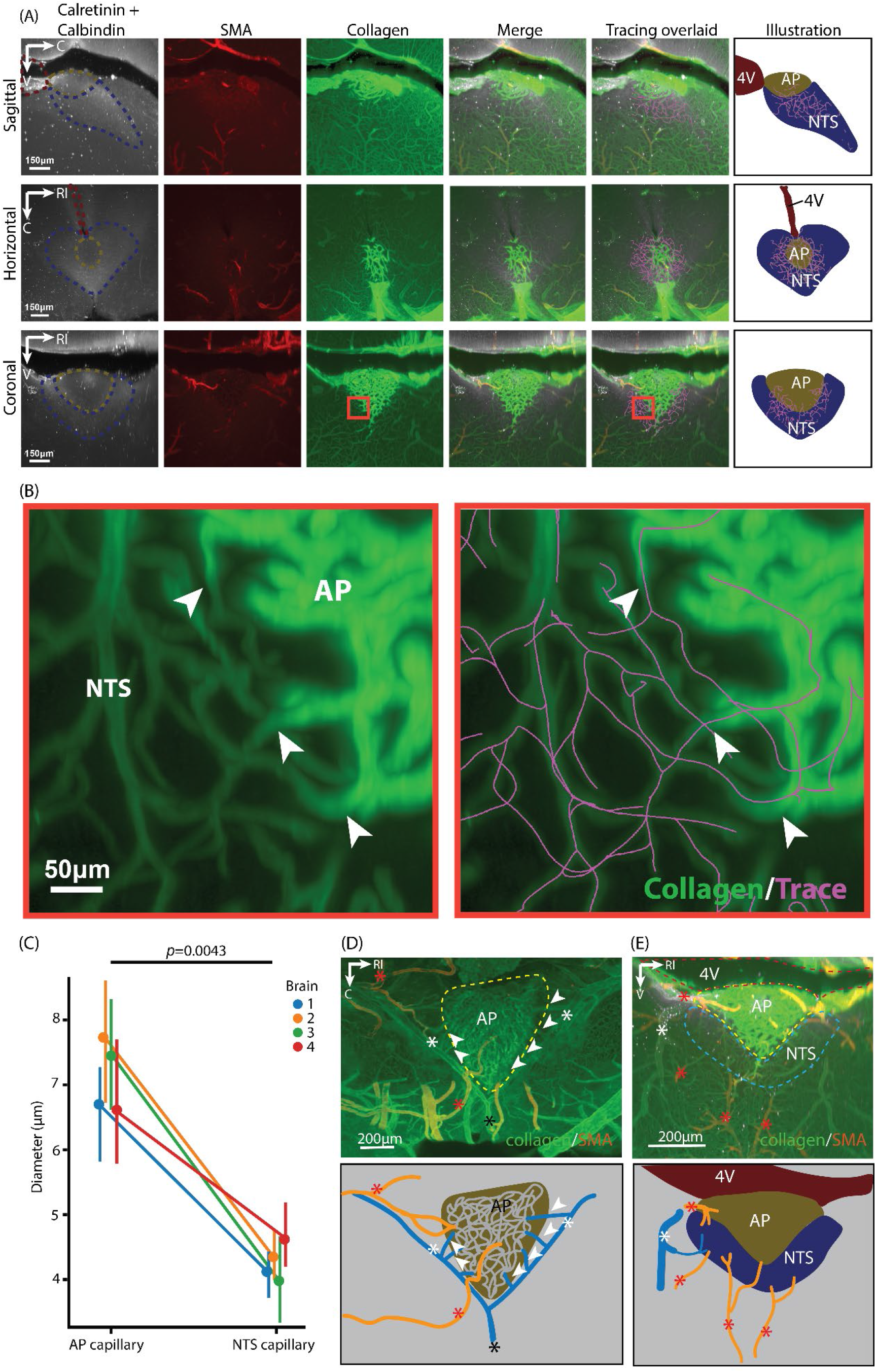
Vascular Connections between AP and NTS in calretinin/calbindin labeled brain slices. (A) Vasculature of AP and the NTS region in sagittal (upper row), horizontal (middle row), coronal (lower row) orientations. For all rows from left to right, NTS labelled by calretinin/calbindin, arteries labelled by SMA, and vasculature labelled by collagen, merge of calretinin/calbindin, SMA, and collagen, merge image overlaid with blood vessel tracing, and their illustration. The ventral, lateral, and caudal surfaces of the AP are covered by the NTS, and the rostral corner of AP is protruding into the 4V. In all three orientations, the results indicate that the capillaries beds of AP and NTS are shared. Color coding as in Figure 4. (B) Magnified view of the boxed region in (A) bottom row 3^rd^ panel (left, collagen staining) and 5^th^ panel (right, collagen merged with tracing). It shows at the border of AP and NTS, the capillaries of both regions form a shared network. Arrowheads point to locations of transitions of capillaries from AP to NTS. (C) Diameter of AP and NTS capillary vessels (4 brains, 5 capillary segments in each region of each brain). The diameter 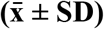 of the AP and NTS capillaries are 7.12±1.15µm and 4.26±0.61 µm respectively. Statistical analysis was performed using repeated measurement ANOVA (F _(1,3)_ = 61.7412, p=0.0043). Detailed statistical results are in Supplementary Table 2. Data in the plot shown as 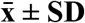. (D) AP arteries and veins in horizontal orientation. Arteries supplying AP reach this area from the caudal and lateral edges (red asterisks). Small vein segments run laterally (arrowheads) from the outer edge of AP, draining to larger veins lying on both sides of AP (white asterisks). These two draining veins converge into a single vein near the caudal corner of the AP (black asterisks). (E) NTS arteries and veins in coronal orientation. The arteries supplying the NTS enter from its lateral and ventral aspects (red asterisks) and the draining veins lie at the lateral aspect (white asterisks). In (A), (B), z=100μm; in (D), (E), z = 300μm. Color code: arteries=orange; capillaries=gray; veins=blue. The remainder of the color code and abbreviations as in Figure 4.

### 3.5 Arteries and veins of AP and NTS

AP and NTS share a joint capillary bed, but they have separate arterial supplies and venous drainage systems. As shown in the horizontal slice (Figure 5C), the artery supplying the AP comes from the lateral sides of medulla and enters the AP from its caudal aspect. AP capillaries merge into a draining vein on each side. The two draining veins converge into a common vein which courses caudally. In the NTS, two supplying arteries can be seen in the coronal view (Figure 5D). One is the penetrating artery from the ventral medulla, entering the NTS from its ventral aspect. The other is the artery traveling along the dorsal surface of the medulla, entering NTS from its dorsal aspect. The NTS is drained by veins converging at its lateral and ventral aspect (Figure 5E).

### 3.6 SFO and AP capillaries connections with the ChP

In addition to their connections with the parenchyma, SFO and AP capillaries also connect to the ChP (Figure 6). In the SFO, capillary branches exit the dorsal SFO and connect to the ChP capillaries (white arrowheads, Figure 6A) that lie near the roof of the 3V, as previously documented^53,54^. The AP capillaries run rostrally and join the ChP capillaries near the caudal 4V, as previously reported^16^.

**Figure 6:**
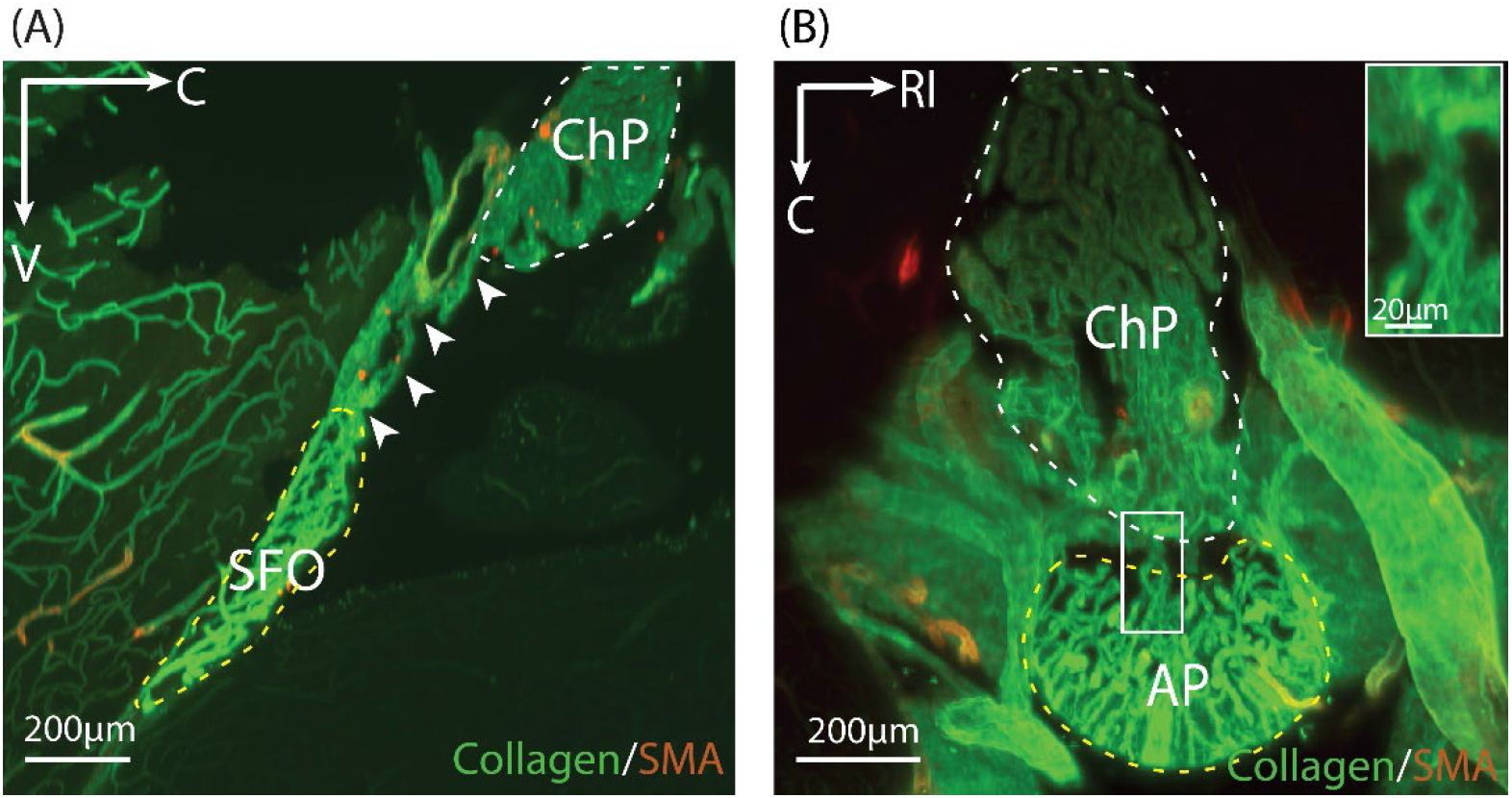
SFO and AP Capillary Connection with the ChP. (A) Capillary from the dorsal SFO connects to the vasculature of the ChP (arrowheads) in sagittal orientation. (B) AP capillaries connected to ChP seen in horizontal orientation. The boxed area is a magnified view of the capillary connection between AP and ChP in the inset. Several capillaries exiting the rostral AP join the capillary vessels of the ChP in the 4V. in Figure 4. In (A) z= 50μm; In (B) z= 300μm. Color coding: dashed outline of AP and SFO = yellow; ChP = white. Abbreviations are as in Figure 5.

## 4 DISCUSSION

### 4.1 Major findings

A portal system linking the SCN and the OVLT has previously been described between the remaining sensory CVO, namely the SCN-OVLT portal system^11^. Shared capillary beds allow diffusible signals to travel from one brain nucleus to another without being diluted in the systemic vascular supply. Such local vascular pathways permit very small populations of neurons to produce secretions that impact specialized local targets. The present work raised the question of whether the other sensory CVOs, namely SFO and AP, share capillary bed connections with the neuropils to which they are attached, as this would provide a means for signals to course directly from one nucleus to the other.

The two major findings in the current study both demonstrate direct vascular connections between the capillary beds of the SFO and AP and the adjacent neuropil. The results indicate the surprising finding of a third portal system in the brain, namely venules joining the capillary beds of the SFO to that of the posterior septal nuclei. This arrangement meets the classical definition of a portal system - an arrangement of two capillary beds bearing vessels are connected by portal veins ^17^. In the other known portal systems in the brain, the capillary bed vessels are of smaller diameter than those of the portal venules^11,55,56^. In the current study, the portal vessels connecting the SFO and the posterior septal nuclei are also larger than the capillaries in both regions, consistent with findings in other identified portal systems (Figure 3C). The second major finding in the current study is that the capillary vessels of the AP connect directly to the capillary bed of NTS with no intervening portal veins. The finding that all three sensory CVOs in the mouse brain bear direct capillary connections to their adjacent parenchyma indicates a new route for local transmission of diffusible signals and a previously unknown pathway for volume transmission.

### 4.2 Relationship of present result to prior literature

Given the tremendous amount of research that has been done on the brain, it is surprising to find new anatomical pathways. Previous research on the whole brain vasculature often failed to report all or some CVOs^57-61^. For example, the BV mapping and analysis in different brain regions by Di Giovanna et al.^57^, Miyawaki et al.^60^ and Ji et al.^58^ was based on high-level organization such as cortex, cerebellum, hippocampus, hypothalamus, thalamus and striatum; Todorov et al.^61^ used finer divisions but didn’t report analysis results in CVOs; Kirst et al.^59^ reported analysis on sensory CVOs but not secretory CVOs. Brain vascular transcriptome studies have deciphered differentiated gene expression in various vascular components^62,63^, but to our knowledge, there are no reports describing the fenestrated capillaries of the CVOs. Our studies of the capillary vasculature of sensory CVOs fill these gaps.

For the SFO, the vascular organization seen in our iDISCO material is in good general agreement with the results of prior work on rats and mice. Thus, the primary arterial supply of the SFO is a branch of the cerebral artery entering the SFO from its caudal direction and the SFO is drained by laterally lying septal veins^64^. Densely tangled capillaries intervene between the arterial supply and venous drainage^65^. Our methods extended this early work by enabling the exploration of vascular connections to adjacent neuropil.

The overall vascular organization of AP found in the iDISCO material is also consistent with previous literature. The primary source of AP arteries is the posterior inferior cerebellar artery, which enters the AP from its caudal aspect^13^. The draining vein of AP described by Kroidl in 1968^66^ is a “venous semicircle” localized to the caudal portion of 4V, circling the posterior half of the AP, which is also seen in the current study. It finally drains to the basilar vein lying at the ventral medulla. We examined AP’s vascular connection with adjacent medulla at a more detailed level however, and here we report findings that differ from the older literature. Roth and Yamamoto^13^ reported that the AP and NTS capillaries are joined by connecting portal veins. Their preparation used coronal brain sections and the “connecting vessels” suggested by Roth and Yamamoto are based on sliced sections which obscures the continuity of the BVs in certain orientations. In our 3D reconstruction of vasculature of this region in the iDISCO material, it is clear that the capillaries of the AP and NTS are interconnected without forming a portal vessel linking the two regions: The capillaries of the two regions are joined, forming a web-like structure seen by viewing the material in various orientations.

### 4.3 Characteristics of vasculature of CVO and their parenchymal attachment sites

It is interesting to note differences in the diameters of capillary vessels in SFO, AP, their portal veins and nearby parenchyma. The general pattern indicates that the portal veins are larger than parenchymal capillaries, and the capillaries in the sensory CVOs are larger than those in the adjacent parenchyma. The difference between the CVO capillaries and their neighboring parenchymal regions is consistent with previous studies, suggesting distinct hemodynamics within different regions^67^. For example, the SFO has longer blood dwelling time than its adjacent parenchyma in the capillary lumen and high blood-to-brain flux rate^68,69^. This is evidenced by comparing the measurements of the amount of radioactive material labeled plasm albumin and diffusible dyes. Albumin in the SFO takes 7-12 times more time to pass through the SFO capillaries compared to blood-brain barrier (BBB) covered capillaries. The radioactive dye is 100-400 times faster to travel across capillary wall compared to the BBB covered capillaries. Another example is that AP microcirculation has significantly larger volume but lower speed of the blood flow compared to other regions in the medulla^68^. It is caused by significantly higher resistance in AP capillaries compared to surrounding parenchymal regions^69^. Gross^68^ hypothesized that these properties allow AP to have enough time to concentrate the blood borne signals traveling to its capillaries, and for transport to its adjacent medulla.

### 4.4 The subregions of SFO and AP have distinct organization

The sensory CVOs have a distinctive vascular organization compared to surrounding neuropil. Unlike other neuropils, they feature an extensive labyrinth of fenestrated capillaries and large perivascular spaces, features assumed to facilitate communication between the nuclei and their humoral milieu (reviewed in^6^). Within the CVOs, the capillary properties are not homogeneous which gives rise to distinct nuclear subdivisions. The SFO has a ventromedial core which abuts the 3V and a shell region neighboring its rostral parenchyma. The SFO core has larger perivascular spaces than the shell^70,71^. In the most rostral part of the shell, there are no perivascular spaces and no fenestrated capillaries^68^. The AP is divided into a mantle region that lies in direct contact with the 4V, a central zone that is the innermost part of AP and lies directly underneath the mantle region, and a ventral zone that surrounds the medial zone and abuts the adjacent medulla (reviewed in ^15^). Unlike the mantle and central zones which bear fenestrated capillaries, the capillaries of the ventral zone are not fenestrated. In addition, the former two zones have clear double-wall capillaries featuring large perivascular space, while the ventral zone shows incomplete fusion of the two walls^72^.

These subdivisional differences indicate that there are some common features of the capillaries in the region forming portal vessels in the SFO and AP. Specifically, there are specialized characteristics of transition zones between the CVOs and their neighboring parenchyma. These zones have little or no perivascular spaces. Also, the capillaries in these zones are less fenestrated than in other subdivisions of the CVOs^67,71-74^. This suggests that the transition zones play a lesser role in transvascular transportation of blood-borne signals and rather participate in the regulation of intravascular movement of diffusible signals.

### 4.5 Implications of joined capillary beds

The discovery of the SFO-posterior septal region portal system and the shared capillary bed of AP and NTS produces a wealth of new research questions, many of which were laid out in the decades that followed the anatomical discovery of a pituitary portal system in 1930^18^,1933^19^ (also reviewed in ^75^). These include determination of signals that flow in the joined capillary beds, the direction and regulation of blood flow, the function of the diffusing signals and mechanism of action at the target sites, etc. Given the unique properties of sensory circumventricular nuclei, including their fenestrated blood vessels and enlarged perivascular spaces, it will be important to explore how they contribute to or participate in the glymphatic systems of the brain^4^. The answers are vital in understanding the functions and the properties of signals that can be transported from one region to the other within each system.

Knowledge of the possible relationship between SFO and the posterior septal nucleus region is limited. To our knowledge, there is no conclusive evidence of neural connections between SFO and TRS or SFi, and they don’t appear to have related functions^76-78^. The SFO is best known for its role in regulating water balance, while SFi and TRS regulate anxiety behavior^79-82^. On the other hand, the AP and NTS are closely related in their functions. As integral parts of the dorsal vagal complex, they regulate cardiovascular functions, feeding behavior and energy balance by integrating peripheral visceral neural inputs, diffusible signals, and afferents from several brain regions^83,84^. The AP seems to be upstream to the NTS, as it sends efferents to the NTS^85^ and possibly diffusible signals travel in the same direction.

### 4.6 Potential significance of CVO vascular organization

CVOs are conservative structures among vertebrate species^86^. They have morphological and cellular differences among different species, but are thought to have similar structure, organization and primary functions^14,16^. Their close apposition to ventricles suggests that they have an ancient origin. Only recently have CVOs been recognized as an indispensable partner in integrating environmental cues and neural inputs for regulating behaviors^87^. Previous findings suggest that communications among brain regions can be achieved by dual pathway system, diffusible routes and neural connections^10,31^.

Together with previous findings of hypothalamic-pituitary portal axis, the current findings of the SFO-posterior septal nuclei portal system and the shared capillary bed of AP and NTS, suggest that connected capillary networks between morphologically distinct nuclei are a fundamental property of CNS organization. Finally, it is noteworthy that the capillary beds of the SFO and AP are also directly connected to the choroid plexus (Figure 6; also reported in ^16,33,53^). Restated, either directly connected or indirectly linked by choroid plexus, the sensory CVOs are all joined together within a single diffusible network with the potential of sharing signaling information between CSF, blood and parenchyma for systemic regulation.

## Acknowledgement

This work is supported by NSF grant 1749500 and NIH grant R21NS134228 (to R.S.). NIH grant DP2MH119423 (to R.T.). Imaging was performed with support from the Zuckerman Institute’s Cellular Imaging platform, and the National Institute of Health (NIH 1S10OD023587-01).

